# Selective attention in hypothesis-driven data analysis

**DOI:** 10.1101/2020.07.30.228916

**Authors:** Itai Yanai, Martin Lercher

**Affiliations:** Institute for Computational Medicine, NYU Langone Health, New York, NY 10016, USA; Institute for Computer Science & Department of Biology, Heinrich Heine University, 40225 Düsseldorf, Germany

## Abstract

When analyzing the results of an experiment, the mental focus on a specific hypothesis might prevent the exploration of other aspects of the data, effectively blinding one to new ideas. To test this notion, we performed an experiment in which we asked undergraduate students to analyze a fictitious dataset. In addition to being asked what they could conclude from the dataset, half of the students were asked to also test specific hypotheses. In line with our notion, students in the hypothesis-free group were almost 5 times more likely to observe an image of a gorilla when simply plotting the data, a proxy for an initial step towards data analysis. If these findings are representative also of scientific research as a whole, they warrant concern about the current emphasis on hypothesis-driven research, especially in the context of information-rich datasets such as those now routinely created in the biological sciences. Our work provides evidence for a link between the psychological effect of selective attention and hypothesis-driven data analysis, and suggests a hidden cost to having a hypothesis when analyzing a dataset.

## Introduction

The pursuit of science may be partitioned into two distinct modes of activity. The great scientist François Jacob had two terms that may better capture the full scientific process, distinguishing between ‘day science’ and ‘night science’ [1]. Day science is the one you read about in the news, it is the one we learn about in school, the one captured by the phrase ‘hypothesis driven’. In contrast, night science is where we are thinking creatively and abstractly, ultimately stumbling upon a new question or a novel hypothesis. And yet when we talk about science, we make it sound as though it is a march of pure rationality, where scientists go from one logical step to another. We have recently developed these ideas in editorials on the creative process of night science [2–5].

Inattentional blindness is a well-known psychological phenomenon that occurs when a person fails to observe an object due to selective attention rather than an inability to observe [6,7]. The best-known experiment of selective attention is called the “invisible gorilla experiment”[8]. In this experiment, volunteers are asked to watch a one-minute video in which six people - three in white shirts and three in black shirts - pass basketballs around. Volunteers are then asked to count the number of passes made by the team in white shirts. While subjects are watching and counting, a person dressed up as a gorilla enters the foreground. Before exiting to the opposite side of the frame, the gorilla pauses, pounding its chest with its fists. While intuition leads us to believe that all of us would see the gorilla, in fact half of the viewers are so focused on counting passes that they completely miss it.

We reasoned that a similar phenomenon may occur when we analyze a dataset; hypothesizing that the mental focus of having a specific hypothesis in mind (analogous to counting basketball passes) would prevent discovery-making in the data.

## Methods

To test this hypothesis, we artificially created a dataset that consisted of 1768 “observations” split across two files labeled “male” and “female”. Each file contained three columns: an ID (ranging from 1 to 1786), distributed randomly across the two files; a column labeled “bmi”, with values between 15 and 32; and a column labeled “steps”, with values between 0 and 15,000.

The dataset was created starting from an image of a gorilla from Classroom Clipart. (https://classroomclipart.com/clipart-view/Clipart/Black_and_White_Clipart/Animals/gorilla-waving-cartoon-black-white-outline-clipart-914_jpg.htm). Using the ‘getpixel’ function in python, we randomly sampled 1786 points from the image to create the dataset, labeling the two dimensions “bmi” and “steps” (Fig. 1). The data was rescaled in gnu R [9] by setting

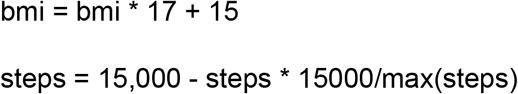

**Figure 1.**
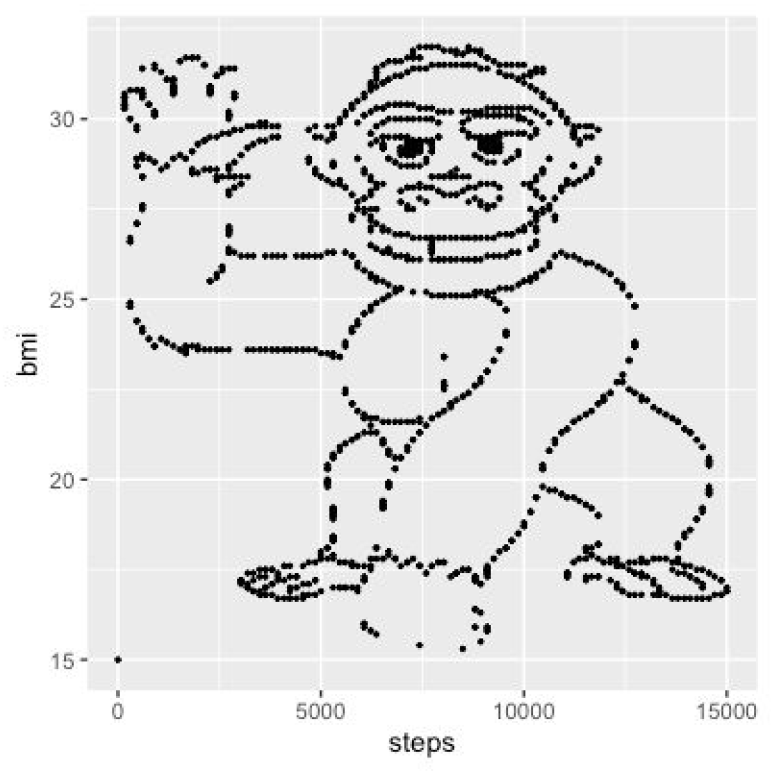
The original gorilla image as a dot plot.

We then split the dataset randomly into two subsets, with an intentional bias towards higher step numbers in the “female” dataset. For this, we first created an index with a random component,

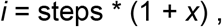

where *x* is a normally distributed random variable with mean 0 and standard deviation 10. We assigned all rows with *i* ≤ median(steps) to the “female” set (*N*_female_=921), and all other rows to the male set (*N*_male_=865).

We provided the two files to 164 undergraduates participating in a course on statistical data analysis for computer science students at Heinrich Heine University Düsseldorf in the context of their weekly assignments. 119 students (here referred to as the *hypothesis-focused* group) received the following instructions as exercise 2 of week 9:

*Download the two files data9b_w.txt and data9b_m.txt. Each row in both files contains for one person (women in data9b_w.txt, men in data9b_m.txt9) the number of steps that this person took on a particular day (steps) and the body mass index (bmi). Assume that both traits are normally distributed for males and for females. Consider the following (alternative, not null) hypotheses:*
  *a)* *There is a difference in the mean number of steps between women and men*.
  *b)* *The correlation coefficient between steps and bmi is negative for women*.
  *c)* *The correlation coefficient between steps and bmi is positive for men*.
  *Think about which test to use and calculate the corresponding P-value.* *Which other conclusions can you draw from the data?*

The remaining 45 students (the *hypothesis-free* group) were provided with an alternative sheet of exercises. Their exercise 2 read:

*Download the two files data9b_w.txt and data9b_m.txt. Each row in both files contains for one person (women in data9b_w.txt, men in data9b_m.txt9) the number of steps that this person took on a particular day (steps) and the body mass index (bmi). Assume that both traits are normally distributed for males and for females.*
*Examine the data appropriately! What do you notice? What conclusions can you draw from the data?*

Note that these instructions are translations, as the original instructions were in German (see SI Text).

Due to the nature of the data, the most notable observation about the dataset was that if you simply plotted the number of steps versus the BMI of either the “male” or the “female” dataset, you would see an image of a gorilla (Fig. 2a,b). We then recorded for each student if they (i) submitted any solutions for the weekly assignment, indicating that they even considered the exercise; (ii) made an attempt at solving the exercise (i.e., went beyond reading the data into their computer); (iii) reported observing the gorilla. One student submitted R code that plotted the data but did not report the gorilla; we excluded this observation, as it is unclear if this person did or did not observe the gorilla.

**Figure 2.**
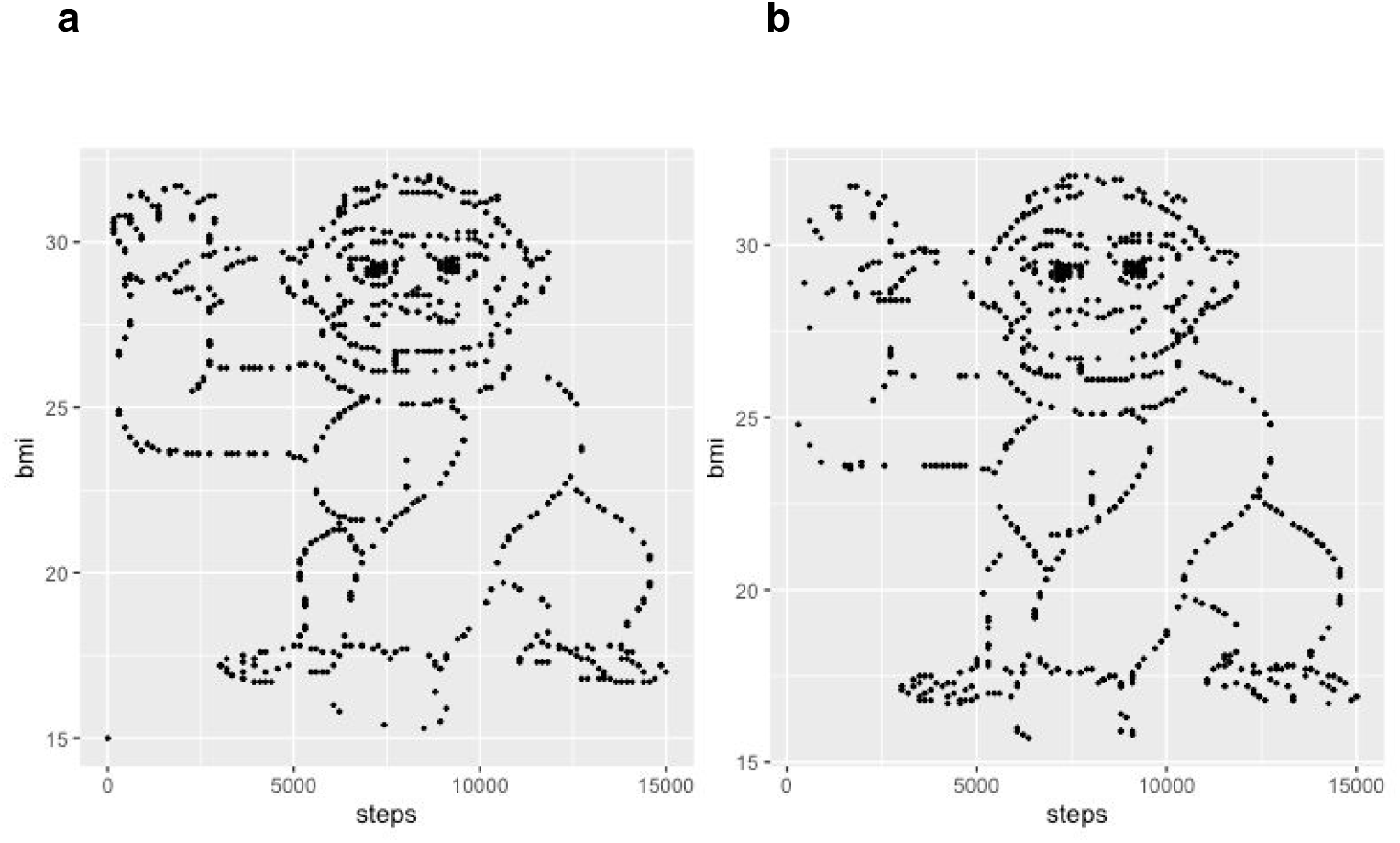
The rescaled gorilla image, randomly split into two biased data subsets. **(a)** “female” dataset; **(b)** “male” dataset.

## Results and Discussion

Table 1 shows the anonymized results for those students who turned in any solutions to the week’s assignment (*N*=44, with *N*_1_=26 in the *hypothesis-focused* group, and *N*_2_=19 in the *hypothesis-free* group). Table 1 summarizes the number of students in each of the two groups who did or did not report observing the gorilla; this table includes only data from students who at least made an attempt at solving the exercise (*N*=33). The odds ratio calculated for the data in Table 1 is 4.80; a one-sided Fisher’s exact test shows that this is statistically significant, *P*=0.034. In other words, students in the hypothesis-free group were almost 5 times more likely to observe the gorilla, our proxy for an initial step towards discovery. Performing the same analysis on all students who submitted solutions to the week’s assignments led to very similar results, with an odds ratio of 4.05 (*P*=0.034, *N*=44).

**Table 1.** Contingency table for reporting the gorilla, summarizing the data in Supplementary Table 1.

|  | Gorilla <u>not</u> reported | Gorilla reported |
| --- | --- | --- |
| Hypothesis-focused | 14 | 5 |
| Hypothesis-free | 5 | 9 |

Table 2 shows a contingency table for all students who handed in solutions to the weekly assignment, contrasting those in the two groups that did or did not attempt a solution that went beyond reading in the data. The odds ratio calculated from this table is 2.15. However, a one-sided Fisher’s exact test shows that this is not statistically significant at the 5% level, *P*=0.21.

**Table 2.**
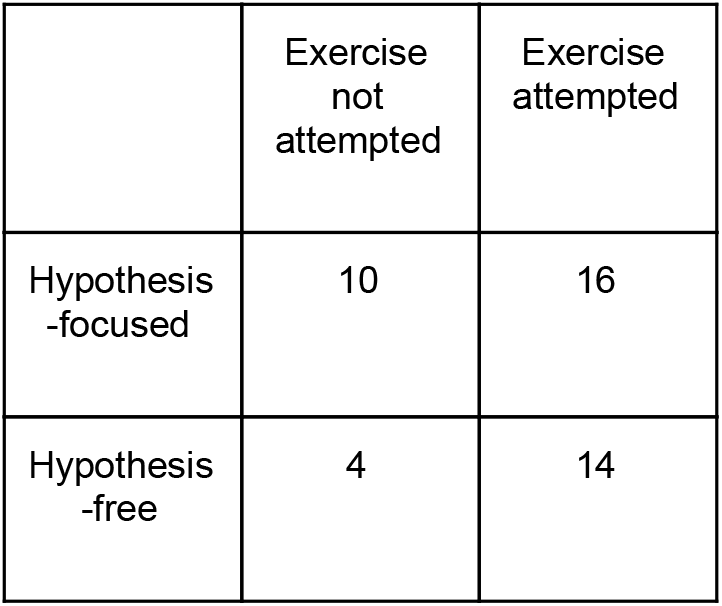
Contingency table for attempting the exercise, summarizing the data in Supplementary Table 1.

|  | Exercise not attempted | Exercise attempted |
| --- | --- | --- |
| Hypothesis-focused | 10 | 16 |
| Hypothesis-free | 4 | 14 |

Thus, we found that a hypothesis-free data analysis is almost five times more likely to make an interesting observation compared to a hypothesis-focused analysis. While we teach our students the benefits of visualization, answering the specific hypothesis-driven questions did not require plotting the data. We found that very often, the students driven by specific hypotheses skipped this simple step towards a broader exploration of the data. Our work thus draws a link between the psychological effect of selective attention and hypothesis-driven data analysis. The results of our small-scale experiment provide evidence that indeed there is a hidden cost to having a hypothesis.

The experiment does have important shortcomings that must be acknowledged. First, we did not control for the time required to perform the tasks across the two conditions, and it may be that the ‘hypothesis-driven’ class was overly tasked and this contributed to ‘missing the gorilla’. While the original selective attention experiment lasted only one minute, for data analysis the temporal aspect is more difficult to control. Second, the specific tasks provided to the participants in the ‘hypothesis-driven’ category are not truly hypotheses, but rather specific tests to perform. We argue though that these are a fair proxy for hypotheses, since these lead precisely to such specific tests.

Presumably, an analyst in a more formal setting may be more attentive to the dataset, particularly if effort was exerted in its generation. While this may be true, we argue that our simple experiment does support the notion that a hypothesis may be a liability for any “night science” explorations. The corresponding limitations on our creativity, self-imposed in hypothesis-driven research, are of particular concern in the context of modern biological datasets, which are often vast and likely to contain hints at multiple distinct and potentially exciting discoveries.

## Acknowledgements

We thank the students of the Statistical Data Analysis course in Heinrich Heine University Düsseldorf for their participation in the described assignment.

## Supplemental Text: German instructions to the students

### Hypothesis-focused group

*Laden Sie die beiden Dateien data9b_w.txt und data9b_m.txt von ILIAS herunter. Jede Zeile in beiden Dateien enthält zu einer Person (Frauen in data9b_w.txt, Männer in data9b_m.txt) die Anzahl Schritte, die die Person an einem bestimmten Tag gemacht hat (steps) sowie den body-mass-index (BMI). Nehmen Sie an, dass beide Merkmale bei Männern und bei Frauen jeweils normalverteilt sind. Betrachten Sie die folgenden (alternativen, nicht Null-) Hypothesen:*
  *a)* *Es gibt einen Unterschied zwischen der durchschnittlichen Schrittzahl von Frauen und Männern*.
  *b)* *Der Korrelationskoeffizient zwischen steps und BMI bei Frauen ist negativ*.
  *c)* *Der Korrelationskoeffizient zwischen steps und BMI bei Männern ist positiv.*
  *Überlegen Sie, welchen Test Sie jeweils verwenden können und berechnen Sie den entsprechenden p-Wert.* *Welche weiteren Schlussfolgerungen können Sie aus den Daten ziehen?*

### Hypothesis-free group

*Laden Sie die beiden Dateien data9b_w.txt und data9b_m.txt von ILIAS herunter. Jede Zeile in beiden Dateien enthält zu einer Person (Frauen in data9b_w.txt, Männer in data9b_m.txt) die Anzahl Schritte, die die Person an einem bestimmten Tag gemacht hat (steps) sowie den body-mass-index (BMI). Nehmen Sie an, dass beide Merkmale bei Männern und bei Frauen jeweils normalverteilt sind.*
*Untersuchen Sie die Daten geeignet! Was fällt Ihnen auf? Welche Schlussfolgerungen können Sie aus den Daten ziehen?*

**SI Table 1.**
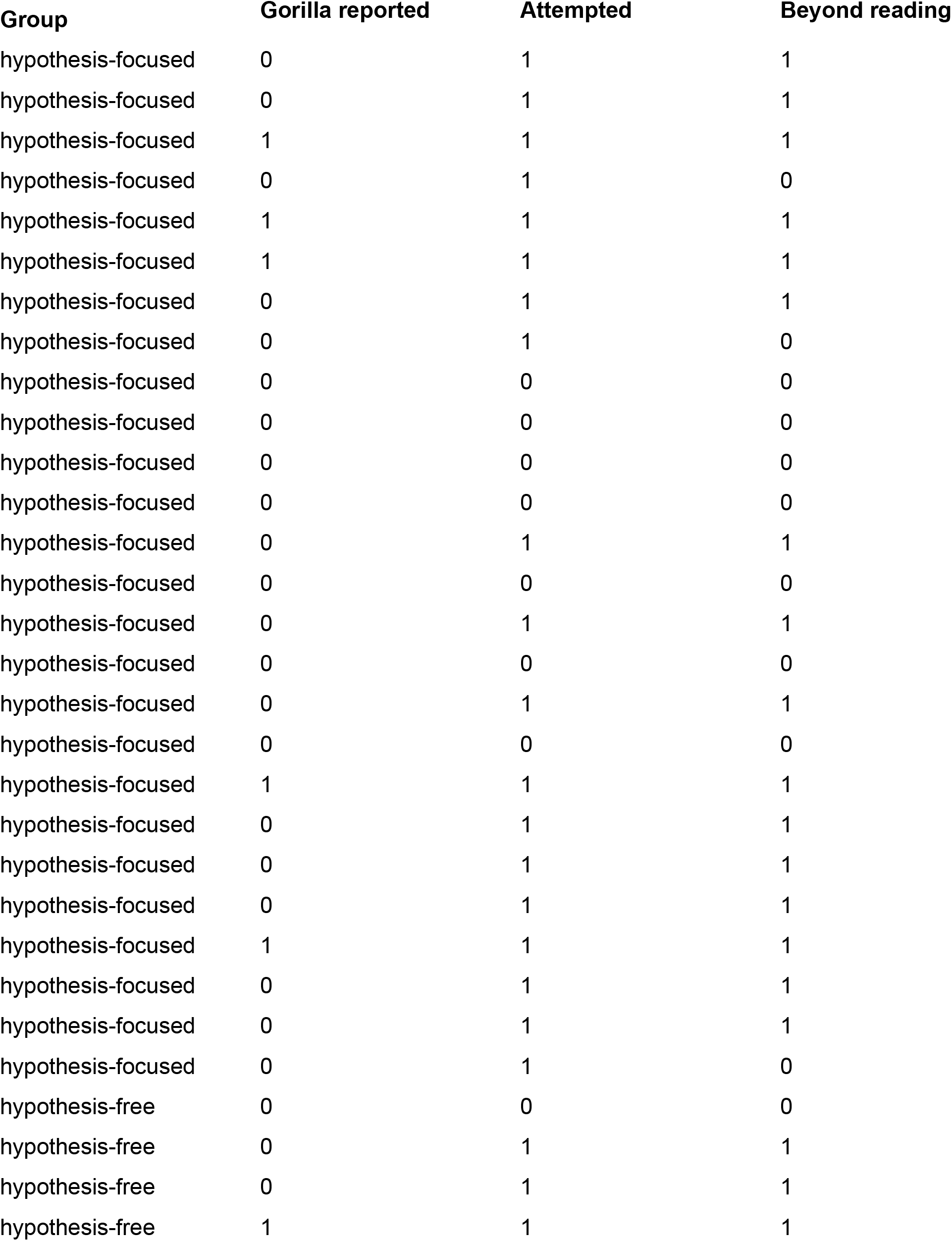
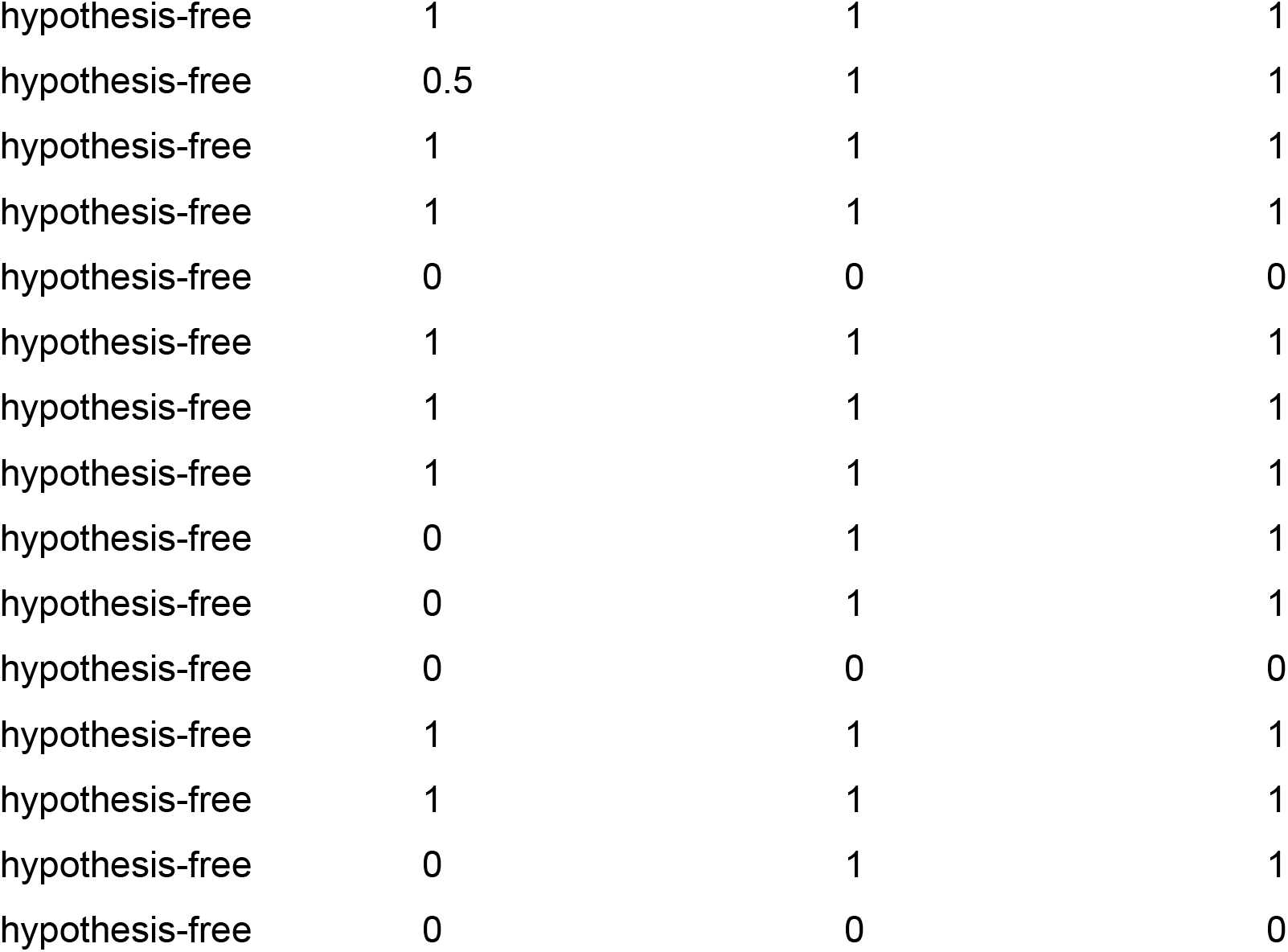
Anonymized data for individual students that handed in solutions to the weekly assignment.

